# Strong transgenerational effects but no genetic adaptation in zooplankton 24 years after an abrupt +10°C climate change

**DOI:** 10.1101/2021.02.05.429921

**Authors:** Antónia Juliana Pais-Costa, Eva J. P. Lievens, Stella Redón, Marta I. Sánchez, Roula Jabbour-Zahab, Pauline Joncour, Nguyen Van Hoa, Gilbert Van Stappen, Thomas Lenormand

## Abstract

The climate is currently warming fast, threatening biodiversity all over the globe. Adaptation is often rapid when the environment changes quickly, but for climate warming very little evidence is available. Here, we investigate the pattern of adaptation to an extreme +10°C climate change in the wild, following the introduction of brine shrimp *Artemia franciscana* from San Francisco Bay, USA, to Vinh Chau saltern in Vietnam. We use a resurrection ecology approach, hatching diapause eggs from the ancestral population and the introduced population after 13 and 24 years (resp. ~54 and ~100 generations). In a series of coordinated experiments, we determined whether the introduced *Artemia* show increased tolerance to higher temperatures, and the extent to which genetic adaptation, developmental plasticity, transgenerational effects, and local microbiome differences contributed to this tolerance. We find that introduced brine shrimp do show increased phenotypic tolerance to warming. Yet strikingly, these changes do not have an additive genetic component, are not caused by mitochondrial genetic variation, and are not caused by epigenetic marks set by adult parents exposed to warming. Further, we do not find any developmental plasticity in response to warming, nor any protective effect of heat-tolerant local microbiota. We conclude that the evolution of shrimp’s extreme thermal tolerance is only due to transgenerational (great)grandparental effects, possibly epigenetic marks set by parents who were exposed to high temperatures as juveniles. This finding challenges standard models of genetic and plastic adaptive responses, and our conception of how species may cope with climate warming.

**Significance statement:** Adaptation is often rapid when environments change quickly, but for climate warming little evidence is available. Many studies report no genetic responses due to pre-existing plasticity, while others point towards epigenetics and microbiota effects. In this study, we take advantage of a natural experiment to study all of these effects. We use a set of coordinated experiments and a ‘resurrection ecology’ approach, reviving resting eggs of brine shrimp up to100 generations after their introduction from a temperate to a tropical saltern. We find that heat adaptation occurs, but heritability is fully “missing”. Plasticity and microbiota play no role either, indicating that only transgenerational (great)grandmaternal effects are involved. This finding prompts us to reconsider the relative importance of the different possible mechanisms by which phenotypic change can occur, especially in response to temperature variation.

## Introduction

Understanding how biodiversity responds to global warming, and anticipating whether species will be able to adapt quickly enough to keep pace with the projected changes, have become major scientific challenges (Hoffmann and Sgrò, 2011). While rapid genetic adaptation to novel human-made environmental changes—polution, pesticides, antibiotics— has been extensively documented (Hendry et al., 2017), much less has been observed for climate warming (Franks and Hoffmann, 2012; Gienapp et al., 2008; Hoffmann and Sgrò, 2011; Merilä and Hendry, 2014; Stoks et al., 2014). This discrepancy might be due to the (yet) modest climate change or to the fact that many pre-existing mechanisms are already in place in most species to cope with the current range of climatic variation.

Theoretically, several mechanisms may cause a phenotypic response to climate warming (Franks and Hoffmann, 2012; Gienapp et al., 2008). First, organisms may genetically evolve to better tolerate high temperatures, and this process may extend their tolerance outside their current thermal niche. They may also phenotypically adjust to these changes using pre-existing plastic responses, within (Chevin and Hoffmann, 2017; Lande, 2015) or across generations (maternal effects, transgenerational epigenetic effects (Auge et al., 2017; Lind and Spagopoulou, 2018)). Finally they may also benefit from symbionts / microbiota adapted to these new conditions (Frankel-Bricker et al., 2020; Nougué et al., 2015; Vannier et al., 2015), without adapting to these conditions themselves. These sources of variation are mutually non-exclusive and can interact in ways that are difficult to disentangle (for instance maternal effects may be mediated by transmitted symbionts, epigenetic marks, or maternal plastic responses (Palumbi et al., 2014; Schlichting and Wund, 2014; Vannier et al., 2015)).

## Results

We investigated whether species can adapt in the wild beyond their climatic niche, with the aim of disentangling these different effects. We used a resurrection ecology approach to assess the thermal adaptive potential of natural populations of the brine shrimp *Artemia franciscana* over 24 years (ca. 100 generations) following an abrupt climatic shift (Lenormand et al., 2018). In the early 1980s, *A. franciscana* from San Francisco Bay, USA (hereafter SFB) were introduced into Vinh Chau salterns, Vietnam (hereafter VCH), where mean (air) temperatures are +10°C higher (Clegg et al., 2000; Frankenberg et al., 2000). This far exceeds the worst IPCC climate warming scenario for the 21st century (RCP8.5 Model predicts +6°C (IPCC, 2013)), yet the brine shrimp have thrived (Van Hoa, 2014), and show phenotypic adaptation to high temperatures (Clegg et al., 2000; Kappas et al., 2004). Indeed, VCH *Artemia* are now commonly used to inoculate other (sub)tropical salterns. We used a series of coordinated experiments to determine the extent to which the introduced *Artemia*’s phenotypic adaptation to higher temperatures resulted from genetic changes, pre-existing plastic responses, transgenerational effects or the effect of locally adapted microbiota (Fig. S1 presents expectations).

We compared the temperature tolerance of an ancestral population from SFB (cysts collected in 1984; hereafter SFB_84_) with that of two populations from VCH (cysts collected in 1997 and 2008; hereafter VCH_97_ and VCH_08_). We resurrected an F0 generation from each population and kept them at a standardized lab temperature, thus removing plastic maternal effects. We then measured juvenile survival in the F1 generation in common garden experiments under temperatures mimicking daily thermal conditions in SFB and VCH (hereafter T_SFB_ and T_VCH_). This experiment was repeated several times as the ‘control’ treatment in the juvenile acclimation, parental acclimation, and microbiota experiments (see below). Very consistently in these controls, VCH populations raised in the laboratory showed increased juvenile survival compared to the original SFB_84_ population, but only when exposed to a VCH climate (meta-analysis χ^2^(1)=9.6, *p*=0.002 at T_VCH_ and χ^2^(1)=0.8, *p*=0.38 at T_SFB_; Fig. 1A solid points). The VCH populations are thus phenotypically adapted to high temperatures, consistent with previous studies (Clegg et al., 2000; Frankenberg et al., 2000; Kappas et al., 2004), and this is not due to direct plastic maternal effects since all F0 females were raised in the same conditions or to different resource allocation of VCH females to their offspring as the effect is specific to VCH temperature. Furthermore, VCH_08_ juveniles had significantly higher survival at T_VCH_ than VCH_97_ juveniles (post-hoc *z*=3.1, *p*=0.002; Fig. 1A), so phenotypic adaptation increased over time in VCH.

**Figure 1.**
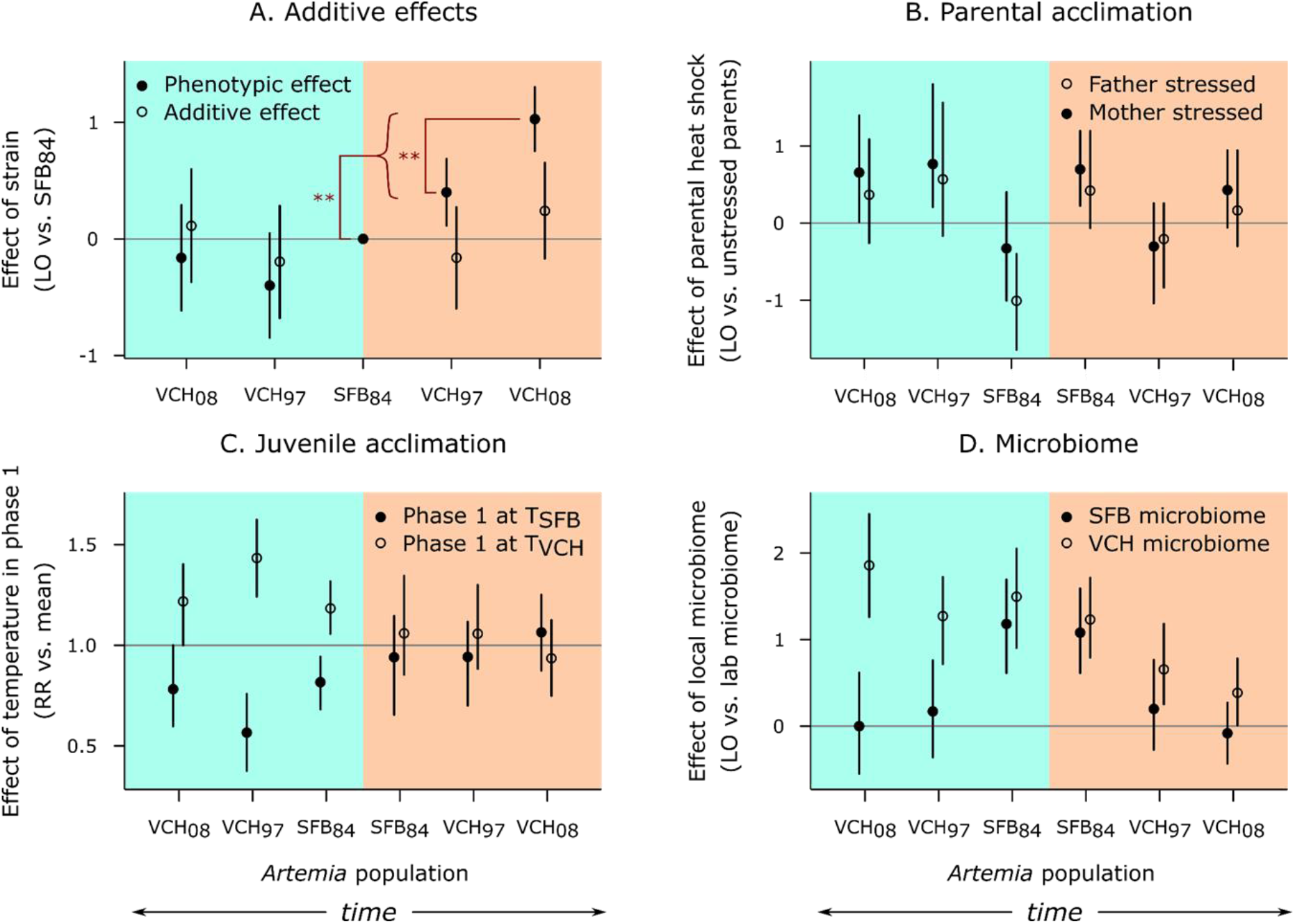
Disentangling the effects of genetics, parental acclimation, juvenile acclimation, and microbiome on phenotypic adaptation to high temperatures. Blue and orange backgrounds represent assays run at T_SFB_ and T_VCH_, respectively. The grey line corresponds to a lack of effect; bars are CIs. To maintain clarity, only significant differences relevant to the phenotypic adaptation to high temperature in VCH are shown; for other *p*-values see Supp. Table 2. Abbreviations: LO, log odds ratio of survival; RR, relative risk of survival. A) Survival of the VCH strains compared to the ancestral SFB_84_, when mothers belonged to the own population (solid points) and to an SFB reference population (‘crossed’ populations, empty points). The ‘0’ point for SFB_84_ is included for reference. B) Difference in survival between the second and first clutches, when parents were exposed to high temperature between clutches 1 and 2. The effect of the second clutch itself (which may have differed in survival compared to the first) is controlled for using the second vs. first clutch effect observed for the unexposed control parents. C) Survival in Phase 2, after exposure to T_SFB_ or T_VCH_ in Phase 1. Here, ‘mean’ is the mean survival in Phase 2 for each strain. D) Survival after inoculation with a local microbiome, compared to survival with the reference lab microbiome.

A second, crucial, step was to determine whether this increased performance resulted from genetic changes. If so, VCH_97_ and VCH_08_ males should be able to transmit at least part of this increased performance to their progeny when crossed with reference SFB females from a stock cultured for over two years under standardized experimental conditions (see methods). This cross removed any maternal and (great)-grandmaternal effects that might have contributed to the observed phenotypic variation. Assuming that adaptation to warmer climate is a polygenic trait, we expect roughly half of the additive genetic effects to be transmitted through males. We would therefore expect to see increased performance at T_VCH_ for the crossed VCH_97_ and VCH_08_ populations but not for the crossed SFB_84_ population in the same juvenile survival test. Despite the strong phenotypic change observed in the uncrossed F1s, survival was not significantly different in juveniles from crossed SFB_84_, VCH_97_ and VCH_08_ populations in either temperature treatment (*p*=0.44 for a population-level difference at T_SFB_; *p*=0.16 at T_VCH_; Table S2, Fig. 1A open points). This means that the increased performance of VCH *Artemia* at T_VCH_ did not result from additive genetic effects. Instead, it may have resulted from (i) fully recessive genetic effects, (ii) maternal genetic effect, notably through mitochondrial evolution, or (iii) plastic grandmaternal (or earlier great-grandmaternal, etc.) effects, e.g. the transmission of epigenetic marks acquired in VCH. Although not formally excluded, the recessivity hypothesis is unlikely as thermal adaptation likely involves numerous quantitative traits with at least some additive effects, and as recessive beneficial alleles (here conferring thermal tolerance) are not expected to sweep quickly in a large population as would be required to explain the rapid phenotypic change observed here (Fig. S3). We investigated the possibility of mitochondrial evolution by sequencing the mitochondrial genome of 10 individuals from SFB_84_ and VCH_08_, as well as sequencing pooled cysts from VCH collected at eight dates between 1984 and 2008. SNP analyses show that mitotype frequencies were remarkably stable over that period, excluding a role for adaptation via the mitochondrial genome (Fig. 2, methods). Hence, it is most likely that the VCH populations have not adapted genetically to higher temperatures. This finding is surprising, but other studies on adaptation to climate warming have also reported an absence of genetic response (Franks et al., 2014; Gienapp et al., 2008; Merilä and Hendry, 2014). Frankenberg et al. (Frankenberg et al., 2000) also showed that VCH *Artemia* populations (hatched from field cysts collected in 1994) had increased survival at high temperature (compared to SFB cysts collected in 1978), but this increased performance was not apparent in later laboratory generations. Such a finding could result from transgenerational effects, supporting our third hypothesis of plastic (great-)grandmaternal effects. Grandmaternal effects are also supported by the study of Norouzitallab et al. (Norouzitallab et al., 2014), who report transgenerational epigenetic effects on thermal tolerance in laboratory *A. parthenogenetica*, which were transmitted up to the F3 generations.

**Figure 2.**
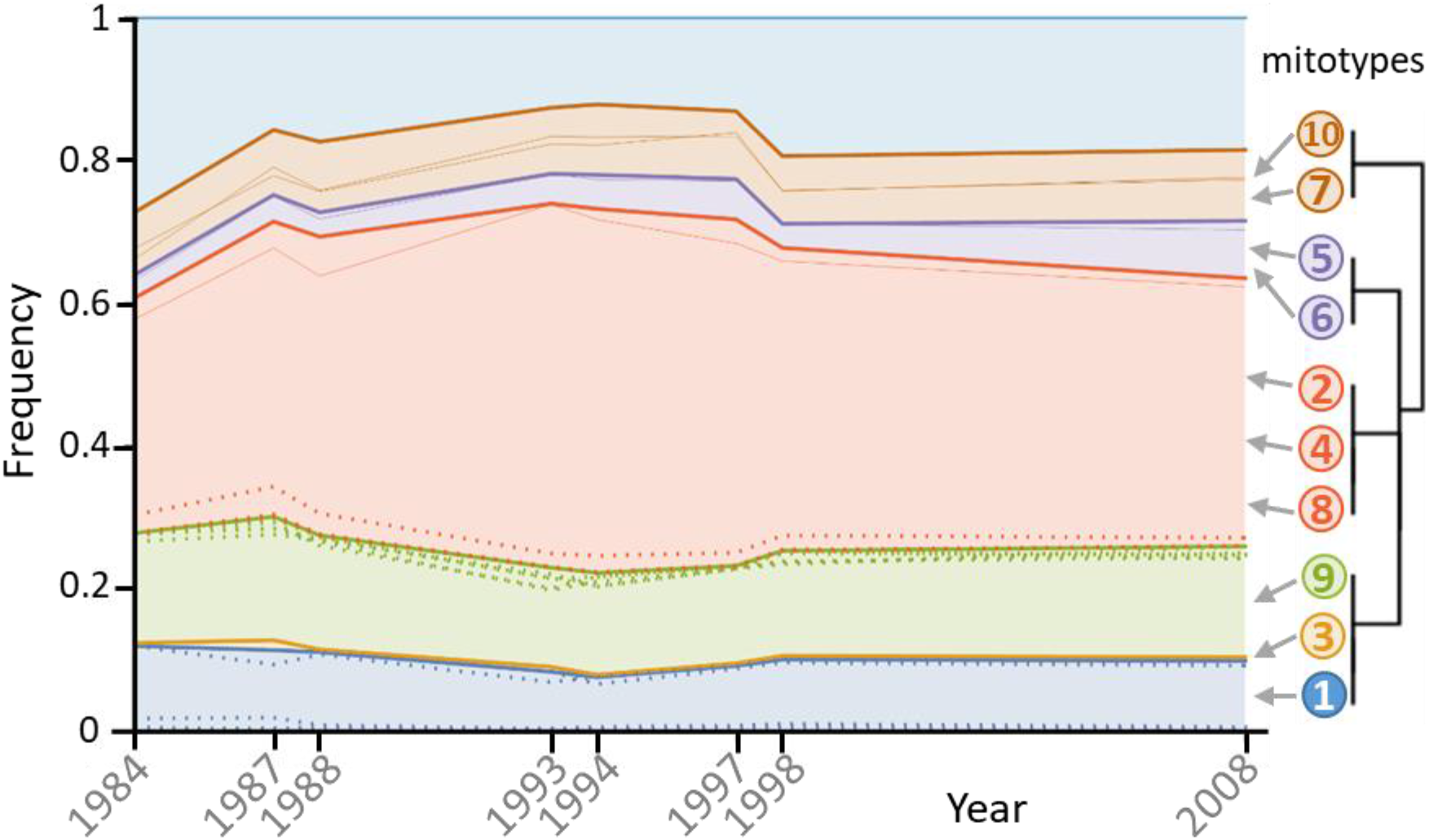
Mitotype frequency variation through time. Sampled years are shown on the *x*-axis, the *y*-axis expresses cumulative frequency. The relationship between the different mitotypes (based on shared-SNP, methods) is shown by the dendrogram on the right. Mitotypes are shown with different colors; numbers identify the individual sequenced (1-5 from 1984 and 6-10 from 2008). The mitotypes’ frequency envelope is that of their most frequent shared-SNP. Individuals 1, 3, and 9 do not have shared-SNPs, and are therefore grouped on this dendrogram. Their frequency envelope is that of their most frequent private-SNP. Thin lines represent other shared-SNP frequencies within mitotypes. Dotted lines represent private-SNPs within groups (only those reaching a frequency >1% are shown).

To further investigate these transgenerational effects, we tested whether thermal exposure of adult parents to T_VCH_ could influence progeny performance at T_SFB_ vs T_VCH_. If so, we would have a mechanism for the grandparental effects (provided they could be maintained for one more generation). We compared juvenile survival in clutches produced before and after exposing their parents to high temperatures (“Parental acclimation” experiment). We exposed the mother, the father, or neither parent. Comparing within the same family controlled for biases resulting from differential mortality of parents exposed to the different treatments; comparisons with families where neither parent was exposed controlled for a second clutch effect. Results showed no significant differences in survival between clutches from the different parental treatments at T_SFB_ or T_VCH_ for any *Artemia* population (0.08≤*p*≤0.36 for a population, parental treatment, or interaction effect at T_SFB_; 0.15≤*p*≤0.41 at T_VCH_; Table S2, Fig. 1B), indicating that thermal exposure in the parent does not influence the thermal tolerance of its progeny. This experiment rules out that epigenetic marks are set in adults in the time window preceding clutch production. However, it does not exclude epigenetic marks that are set during the juvenile development of the parents (or grandparents, etc.) (Donelson et al., 2018; Norouzitallab et al., 2014). The imprint may be set early during meiosis in the female germ line, which occurs during juvenile development (Lenormand et al., 2016). Indeed, the epigenetic effects referenced above were found after exposing juvenile *A. parthenogenetica* to a heat shock (Norouzitallab et al., 2014).

Next, we investigated whether *Artemia* have a developmental plasticity response to the thermal environment. Such a plastic response would not be sufficient to explain the phenotypic effects that we observed, since these experiments did not include an acclimation phase before measurement. However, if plastic adjustment to cope with high temperatures pre-existed in SFB, or evolved in VCH, this would help explain the lack of genetic change in VCH. This possibility is reinforced by previous studies in *Artemia*, which demonstrated plastic response to thermal stress (Clegg et al., 2000; Frankenberg et al., 2000). To investigate this possibility, we exposed 5-day-old juveniles to T_SFB_ or T_VCH_ for 2 days, and then tested whether pre-exposure increased performance in each environment (“Juvenile acclimation” experiment) during the same age window used for the other experiments. Strikingly, we found that early exposure to T_VCH_ did not increase juvenile survival at T_VCH_ in any of the *Artemia* populations (*p*≥0.62 for an effect of pre-exposure or its interaction with population; Table S2, Fig. 1C). In contrast, pre-exposure to T_VCH_ significantly increased survival at T_SFB_ (*p*<0.0001 for a pre-exposure effect; Table S2, Fig. 1C) for all three *Artemia* populations (*p*=0.31 for an interaction with population; Table S2), indicating that there is indeed a plastic response (e.g. activation of heat shock proteins (Clegg et al., 2000; Frankenberg et al., 2000)). However, this plasticity does not confer improved performance at T_VCH_, so we can rule out that it plays a role in the thermal adaptation at VCH.

Last, we investigated whether performance at T_SFB_ and T_VCH_ could be affected by the presence of microbiota adapted to those climates (“Microbiota” experiment). In corals, for example, the temperature niche is controlled by that of their symbionts (Littman et al., 2010). *Artemia* host many gut bacteria that are essential for the proper digestion of unicellular algae, their main food source. Adaptation of this microbiota to high salinity has been shown to determine their host’s salinity niche (Nougué et al., 2015). Hence, it is possible that *Artemia*’s thermal niche is controlled in part by the thermal niche of its microbiome. Such a finding would also help explain the lack of genetic change in VCH. To evaluate this possibility, we investigated the thermal tolerance of axenic *Artemia* from SFB_84_, VCH_97_, and VCH_08_ populations inoculated with microbes sampled from live *Artemia* in SFB, VCH, or our reference laboratory cultures. If microbes contribute to thermal tolerance, we would expect VCH microbes to increase juvenile survival at T_VCH_, but not T_SFB_, whereas SFB microbes should increase survival at T_SFB_ but not T_VCH_ (Fig. S1). We did not find this pattern. Instead, we found that having microbes from VCH increased survival for all *Artemia* populations at both T_SFB_ and T_VCH_, while having lab microbes decreased survival in all circumstances (*p*=0.003 for an interaction between population and microbiome at T_SFB_; *p*=0.001 at T_VCH_; Table S2, Fig. 1D). Hosting VCH microbes appears to simply be better than hosting lab microbes. For the SFB microbes, we found that they conferred the same survival as VCH microbes in SFB_84_ but were equally poor as the lab microbes for both VCH populations. Hence, our results are consistent with the idea that (i) microbes have a large impact on survival, (ii) microbes from our three stocks are different and (iii) their effect depends on the *Artemia* population. We did not find any indication that the microbes play a role in thermal adaptation. Interestingly, we found that *Artemia* had no problems when exposed to microbiota from a tropical climate: they are available, and there is no need to specifically adapt to them (as SFB_84_ performed equally well with VCH microbes). All our findings are consistent with a loss of function in the laboratory microbes, and by a loss of ability of the Vietnamese *Artemia* to cope with their ancestral SFB microbes.

## Discussion

In summary, we found no indication of genetic adaptation to increased temperature in a field situation which should *a priori* be very favourable for the evolution of thermal tolerance (Reznick and Ghalambor, 2001): a large and isolated sexual population without initial bottleneck, exposed to a large and abrupt environmental shift over 100 generations. However, we did find a phenotypic difference when testing individuals whose grandmothers were exposed to high temperatures, and this difference was larger for the VCH_08_ population than for VCH_97_. We conclude that VCH *Artemia* have higher heat tolerance due to transgenerational effects, and that these effects increased through time, e.g. by being better maintained through generations in more recent Vietnamese populations. Such effects are not entirely unexpected, as they are found more often in short‐lived, dispersal‐limited organisms, for juvenile traits, and in conditions where environmental variation is predictable over several generations (Yin et al., 2019). Further work is necessary to see whether these transgenerational effects are due to epigenetic marks, as in (Norouzitallab et al., 2014), or due to other mechanisms. Their presence could explain the lack of genetic changes in VCH: transgenerational effects could keep the population phenotype close to a thermal optimum, thereby reducing directional selection and genetic changes. Such interference with genetic adaptation has been found in many studies reporting within-generation plasticity (Gienapp et al., 2008; Merilä and Hendry, 2014), but the transgenerational mechanism found here is much less documented. Transgenerational effects may be difficult to detect: though the resurrection ecology approach is among the most powerful methods to study adaptation to climate change (Lenormand et al., 2018; Nogués-Bravo et al., 2018; Orsini et al., 2013; Weider et al., 2018), without crosses we would have concluded that genetic adaptation had taken place (as in Geerts et al., 2015; Yousey et al., 2018). Compared to other studies, our adaptive transgenerational effects are large (Jeremias et al., 2018; Sánchez‐Tójar et al., 2020; Yin et al., 2019), and contrast strikingly with the absence of adaptive genetic and within-generation plastic effects, providing an example where adaptation involves traits whose heritability is entirely “missing” (Trerotola et al., 2015). Overall, this work represents one of the most complete studies jointly addressing the different factors associated with thermal adaptation in the wild, namely genetic effects, epigenetic effects, plasticity, and microbiota. It shows that phenotypic adaptation to an extreme environmental change can be achieved by transgenerational effects and suggests that these effects can be so large that no genetic adaptation takes place.

## Acknowledgments

We thank N. Rode, G. Martin, L.-M. Chevin, and A. Charmantier for contributing to a pilot experiment; Y. Michalakis, C. Teplitsky, S. Glémin, L.-M. Chevin and N. Bonel for suggestions and comments on the manuscript; C. Mahieu for maintaining the ARC cyst collection; and Susan de La Cruz and David Nelson from San Francisco Bay Estuary Field Station for providing live SFB Artemia for the microbiota experiment. We thank M.-P. Dubois and The Genomics, Molecular Ecology, and Experimental Evolution platform (GEMEX) at CEFE; the genotyping and sequencing facilities of the Institut des Sciences de l’Evolution (GENSEQ); the Labex Centre Méditerranéen de l’Environnement et de la Biodiversité (CEMEB); and the MGX sequencing platform in Montpellier. Financial support was provided by MUSE Kim Sea and Coast, CNRS and Fundación BBVA (Ref 201632009) and FCT Portugal, through MARE (UID/MAR/04292/2013), and through the PhD grant of A. J. Pais-Costa (SFRH/BD/108224/2015).

## Methods

Experiments were performed with three populations of *A. franciscana*: one from San Francisco Bay (SFB), USA, collected in 1984 (SFB_84_); a second from Vinh Chau (VCH) saltern, Vietnam, collected in 1997 (VCH_97_); and a third, also from VCH, collected in 2008 (VCH_08_).

### Decapsulation and hatching of the cysts

The parental generation of experimental individuals (see below) was hatched from dormant cysts. Cyst decapsulation and hatching protocols were modified from Bengtson et al. (1991). Cysts were rehydrated in deionized water (2 h to 3 h). After rehydration, cysts were decapsulated by a 10 min exposure to a sodium hypochlorite solution, then rinsed with running water (10 min) and deionized water (5 min). Decapsulated cysts were incubated for 48 h at 28 °C (± 1°C), with constant light and aeration, in a 5 g/L salinity medium (see below). After emergence, first-instar nauplii were moved to 23 °C (±1°C) and natural light conditions. Salinity was gradually increased to 80-90 g/L over 8-9 days. This procedure was performed independently for the three different *Artemia* populations.

### Baseline experimental conditions

Throughout the preparation and execution of the experiments, *Artemia* were kept in a 80-90 g/L saline medium prepared by diluting field-collected concentrated brine (280 g/L, Camargue Pêche, France) from Aigues-Mortes saltern with deionized water. Organisms were fed a solution of *Tetraselmis chuii* algae (Fitoplankton marino, Spain), prepared by dissolving 1 g of lyophilized algae in 1 L of deionized water (c.a. 6.8×10^9^ *T. chuii* cells/L). Stock individuals were fed ad libitum. Food was added daily (1ml of algae /per group of juveniles / day; and 1ml of algae / per couple / day) before exposure and 3 times a week (2 ml of algae / group of juveniles / 2 days) during exposure to the temperature treatments. Unless specifically mentioned, individuals were kept at 23 °C (±1 °C), under natural light conditions. Juvenile survival tests were all performed in the dark in incubators and thermostatic chambers. Mortality was checked twice (5 and 10 days after the beginning of the treatment, i.e. midway through and at the end of the thermal treatment).

### SFB and VCH temperature regimes

The same temperature regimes were applied in each experiment. Two temperature cycles were used: i) cycle of temperatures based on the air temperatures from SFB (T_SFB_): 16 °C (2 h); 22 °C (8 h); 27 °C (4 h); 22 °C (8 h); 16 °C (2 h); and ii) cycle of temperatures based on the air temperatures from VCH saltern (T_VCH_): 26 °C (2 h); 32 °C (8 h); 37 °C (4 h) for experiment 2 and 35 ºC (4h) for the remaining experiments; 32 °C (8 h); 26 °C (2 h).

### Experiment 1: microevolution/adaptation

This experiment was performed to measure the additive genetic effect of thermal adaptation, removing maternal lineage effects. For this experiment, we collected virgin females from a laboratory population of *A. franciscana* from SFB, hatched from cysts collected in 2003 (SFB_03_). The SFB_03_ population was maintained in the laboratory for over 2 years, so it was well acclimated to the standard laboratory temperature conditions (23°C ± 1°C). For SFB_84_, VCH_97_, and VCH_08_, we hatched individuals from field cysts. Before individuals reached sexual maturity, their sex was assigned based on sexual dimorphism. After maturity, males from the three study populations of *Artemia* were mass crossed (animals divided into 4 replicates) with the virgin stock females (SFB_03_) to produce an F1 generation. Starting 24h after the first nauplii were seen, we collected daily batches of nauplii from the mass crosses. This ensured that the organisms used in each replicate were born within the same period. Newborn nauplii from each cross were maintained in 50 ml Falcon tubes (max. 30 nauplii per tube) filled with 30 ml of brine solution for a period of 7 days. After 7 days, all meta-nauplii from the same cross were mixed and then separated into replicate groups of 10 individuals. Each group was placed in a 50 ml Falcon tube filled with 30 ml of brine solution, and exposed to T_SFB_ or T_VCH_ for 10 days (7^th^ to 17^th^ day). 30-32 groups per population were exposed to each cycle of temperatures (1830 individuals in total).

### Experiment 2: Parental acclimation

This experiment was designed to investigate the possibility that thermal exposure in the parents could influence juvenile performance at high temperature. Individuals were hatched from SFB_84_, VCH_97_, and VCH_08_ field cysts. Before they reached sexual maturity, their sex was assigned based on sexual dimorphism. After maturity, single pairs of males and females from each population were isolated in 50 ml Falcon tubes filled with 30 ml of brine solution to produce an F1 generation. We collected the first brood of nauplii produced by each parental couple. Each brood of nauplii was isolated from their parents after confirming, under a stereomicroscope, that the female ovisac was empty. In this way, we ensured that the organisms used in each replicate were born within the same period. Immediately after the first clutch (CL_1_) was born, the parents were separated, and one of three treatments was applied: i) mother exposed to high temperature; ii) father exposed to high temperature; and iii) control (none exposed to high temperature). The ‘high temperature’ treatment consisted of 8 h at 35 °C (±1 °C) in the dark. Afterwards, the couples were put back together to produce a second clutch (CL_2_), which we collected in the same way. Newborn nauplii were kept in 50 ml Falcon tubes (max. 30 nauplii per tube) filled with 30 ml of brine solution for a period of 7 days. After 7 days, meta-nauplii from each family were separated into groups of 10 individuals and placed in 50 ml Falcon tubes filled with 30 ml of brine solution. A minimum of 5 replicate groups were tested per couple. Each replicate was exposed to T_SFB_ or T_VCH_ for 10 days (7^th^ to 17^th^ day). Overall, 39.3 groups (SD 9.4) were used per treatment, population, and temperature regime combination (survival of 7080 individuals assayed in total).

### Experiment 3: Juvenile acclimation

This experiment was conducted to study if very early exposure of the organisms to a thermal regime would increase their performance as juveniles under the same regime. Individuals were hatched from SFB_84_, VCH_97_, and VCH_08_ field cysts. Before individuals reached sexual maturity, their sex was assigned based on sexual dimorphism. After maturity, single pairs of males and females from the same population were placed in 50 ml Falcon tubes filled with 30 ml of brine solution to produce an F1 generation. We collected newborn nauplii from the parental couples. Each brood was isolated after confirming, under a stereomicroscope, that the female ovisac was empty. In this way, it was ensured that the organisms used in each replicate were born within the same period. Newborn nauplii were counted and separated into 50 ml tubes containing 30 ml brine solution (maximum 30 nauplii per tube). Nauplii were then maintained under the same conditions of light, food and temperature as the parents for a period of 5 days. After 5 days, meta-nauplii entered the experiment, which was divided into 2 phases (P1 and P2). At day 5, a first temperature regime was applied for two days (P1). Meta-nauplii from each family were separated into 50 ml Falcon tubes (max. 30 nauplii per tube) filled with 30 ml of brine solution and assigned to either T_SFB_ or T_VCH_. Broods were discarded whenever it was impossible to obtain two replicates (one per temperature regime) with a minimum of 10 individuals each. After this first phase (P1), mortality was checked, and surviving meta-nauplii were separated into groups and placed into 50 ml Falcon tubes (no more than 14 individuals per falcon) filled with 30ml of brine solution. Broods were discarded whenever it was impossible to obtain two replicates (one per temperature regime) with a minimum of 5 individuals each. Meta-nauplii from each population and temperature regime were again assigned to T_SFB_ or T_VCH_ for the second phase (P2). Hence, different individuals were exposed to different temperature histories: T_SFB_ → T_SFB_, T_SFB_ → T_VCH_, T_VCH_ → T_SFB_, T_VCH_ → T_VCH_. Survival during this second phase was recorded for a period of 10 days (7^th^ to 17^th^ day). Overall, 48.8 (SD 12.0) groups were used per temperature history (P_1_ → P_2_) and population combination (survival of 5315 individuals assayed in total).

### Experiment 4: Microbiota

This experiment was designed to investigate whether exposing organisms to microbiota adapted to different climates lent their hosts different performance in those climates. SFB_84_, VCH_97_, and VCH_08_ field cysts were rehydrated in sterile deionized water (2 h to 3 h). After rehydration, cysts were decapsulated by a 10 min exposure to a sodium hypochlorite solution, then rinsed with deionized water (10 min) and sterile deionized water (5 min). Decapsulated cysts were then incubated for 3 days at 28 °C (±1 °C) and under constant light, in sealed bottles containing 400 mL autoclaved brine solution (5 g/L). After emergence, first-instar nauplii were placed at 23 °C (±1 °C) under constant light and fed with sterilized *T. chuii* solution. This procedure was performed independently for the three different populations. Salinity was gradually increased to 80-90 g/L over 8-9 days. When salinity reached 40 g/L, nauplii from each population were separated into 3 groups and inoculated with i) microbiota from SFB ii) microbiota from VCH or iii) microbiota from containers in the laboratory. The microbiota inoculum was obtained by mixing crushed live adult individuals collected in 2017 in two sites in both Vinh Chau saltern (salinity 70 g/L and 90 g/L) and San Francisco Bay Estuary Field Station (salinity 70g/L and 130 g/L). These 2017 microbiota might differ from the original 1984 situation, but the thermal background did not significantly changed between 1984 and 2008 and those microbial communities should reflect this climatic difference. Each inoculation bottle was filled with 400 ml of sterile deionized water and 100 ml of microbiota solution. Sterilized *T. chuii* was added ad libitum. When individuals reached sexual maturity, 12 males and 12 females from each population and treatment were separated into new sterile containers and mass crossed to produce a F1 generation and kept under the same conditions as the stock. Newborn nauplii were checked daily. Each batch of nauplii was isolated within 24 h after the first nauplius was seen, to ensure that organisms used in the experiment were born within the same period. Newborn nauplii were separated into sterile 50 ml tubes (max. 30 nauplii per tube) containing 26 ml of sterile brine solution, 2 ml of microbiota solution, and 2 ml of autoclaved algae solution. Nauplii were then maintained under natural light at 23 °C (±1 °C) for a period of 7 days. After 7 days, all meta-nauplii from the same treatment were mixed and separated into replicate groups of 10 individuals. Each group was placed in a sterile 50 ml tube containing 26 ml of sterile brine solution, 2 ml of microbiota solution, and 2 ml of algae solution. To maintain the comparison with the other experiments, only the water was autoclaved to prepare the food solution for the rest of the experiment (i.e. not the lyophilized algae, which would have significantly altered the food source). Each replicate was exposed to T_SFB_ or T_VCH_ for 10 days (7^th^ to 17^th^ day). Overall, 87-103 groups (27-39 groups per microbiota treatment) per population were exposed to each temperature regime (5630 individuals in total). All feeding and transfers were performed under a laminar flow hood to prevent microbial contamination. During the experiment, the containers were closed to limit contamination, but not sealed to allow gas and oxygen exchange.

### Statistical analyses

We first analysed the overall temperature tolerance of the VCH populations compared to the ancestral SFB_84_. To maximize our power to detect differences between populations, we pooled the ‘control’ data from the acclimation and microbiome experiments. Specifically, we used: the first clutches from the ‘Parental acclimation’ experiment, the second clutches from the ‘Parental acclimation’ experiment whose parents were not exposed to high temperature; the individuals from the ‘Microbiome’ experiment who were inoculated with the lab microbiome; and the organisms from the ‘Juvenile acclimation’ experiment who had undergone the same temperature regime in Phases 1 and 2. There are of course some small differences between these experiments (i.e. the ‘Juvenile acclimation’ organisms had undergone a slightly longer exposure to the temperature regimes, the ‘Microbiome’ organisms were cultured differently), but the meta-analysis approach accounts for this additional variation. We used a multilevel meta-analysis model (R Core Team, 2014; Viechtbauer, 2010), and meta-analysed the two temperature regimes separately. Survival relative to the SFB_84_ population was taken as the response variable because it is the ancestral population. Effect sizes were obtained by fitting binomial models like those described below to the control data for each experiment, and extracting the log odds ratio of each VCH population relative to SFB_84_ (more details in Suppl. Table 1). Standard errors extracted from the same models were used to weight the meta-analysis. The full meta-analysis model contained *VCH population* (VCH_97_ or VCH_08_) as a fixed effect, and *Experiment* as a random effect controlling for non-independence within experiments. The significance of *VCH population* was then tested using likelihood ratio tests. Where relevant, post-hoc Tukey tests were performed to compare the two populations.

To analyse the individual experiments, we used generalized linear mixed models (Bates et al., 2015; R Core Team, 2014), with the number of surviving vs. dead *Artemia* in each replicate as the response variable (binomial response with logit link). The two temperature regimes were analysed separately (i.e. the following was repeated for T_SFB_ and T_VCH_). First, we constructed a full model that included all the experimentally manipulated factors and their interactions. The ‘Additive genetic effects’ models included only *Population*. The ‘Parental acclimation’ models included *Population*, *Clutch* (a dummy variable, with the first clutch coded as ‘0’ and the second clutch as ‘1’), and their interaction, and the interactions between these and the factor *Parental treatment*. By using the dummy variable and restricting *Parental treatment* to the interaction terms, we avoided generating spurious (and biologically impossible) estimates of the effect of *Parental treatment* on the first clutch. We also included a random *Family* term to group replicates collected from the same parental couple. In the ‘Juvenile acclimation’ experiment, we analysed the survival in Phase 2, which was conditional upon survival in Phase 1. The models included *Population*, *Temperature in Phase 1*, and their interaction, as well as a random *Family* term to group replicates collected from the same parental couple. For the ‘Microbiome’ experiment, the models included *Population, Microbiome* and their interaction. Where necessary, the full models were corrected for overdispersion by including an observation-level random effect (Harrison, 2015). Finally, the significance of the predictors was tested using likelihood ratio tests. For the ‘Parental acclimation’ experiment, where we were only interested in the effects of *Population* and *Parental treatment* on the difference between the first and second clutch, we only tested the significance of the interaction terms.

### Mitochondrial genome sequencing and analyses

In order to determine whether increased heat tolerance of the Vietnamese populations could be caused by mitochondrial genetic variation, we sequenced the full mitochondrial genome of 10 individuals (individuals 1-5 sampled in 1984, and individuals 6-10 sampled in 2008). We also sequenced pools of cysts sampled in Vinh Chau saltern (25mg of cysts, ca 6500 cysts per pool) from eight years (1984, 1987, 1988, 1993, 1994, 1997, 1998, 2008). Three of these were replicated twice, with independent DNA extraction (1984, 1997, 2008). For each sample, mitochondrial DNA was extracted using an Abcam ab65321 Mitochondrial DNA isolation kit, following the manufacturer’s instructions. NGS libraries were constructed using a Nextera DNA flex illumina kit (ref 20018704) and sequenced (PE 150) on an Illumina NovaSeq 6000 (MGX platform, Montpellier).

For each sample, paired reads were mapped onto an *A. franciscana* reference sequence (NC_001620.1) with *bowtie2*, trimming 10 bases in 5'. Read duplicates were removed with *Picard MarkDuplicates*. Reads with a mapping quality over 20 and in proper pairs were kept with *samtools view*. The program *pysamstats* was used to get the raw percentage of each base and the total coverage at each position of the reference sequence. These steps were done twice, on the original reference genome and on a version that was cut in the middle and had the two parts reordered. This was done to avoid border effects and obtain a good mapping for the reference extremities of this circular genome. A dedicated *R* script was written to concatenate the *pysamstats* output files, keeping 50% middle positions of the two reference versions, to obtain two tables with all samples: one with the percentages of alternative bases at each position and one with the coverages. SNP calling was done using a dedicated Mathematica 10.1 (*Wolfram*) script. Genome coverage was ~3000X on average for cyst pool samples (range 1000X – 6000X), and was ~200X on average for individual samples (range 42X-336X). Three regions showed a drop in coverage on the reference genome and were excluded from further analyses (region 1: 14045-14394; region 2: 14682-14835; region 3 15409-15806).

Forty variable positions were identified that distinguished the 10 sampled individuals. One of them was an ambiguous insertion of a variable number of T’s at position 1247, and was removed. Among the 39 remaining SNPs, 7 were shared by at least two individuals and 32 were private to a single individual. The shared-SNPs defined 6 non-ambiguous haplotypes (hereafter ‘mitotypes’), three being characterized by a combination of at least two shared-SNPs (individuals 7 and 10; individuals 5 and 6; individuals 2, 4, and 8) and three by the absence of shared-SNPs (individuals 1, 3, and 9). The frequency envelopes of the former were obtained using the frequency of their most frequent shared-SNP, while the frequency envelope for the latter was based on the frequency of their most frequent private-SNP (as in the absence of recombination, the sum of the frequency of private SNPs cannot exceed that of shared SNPs within a mitotype).

The frequency of each of the 39 SNPs was estimated from the cyst pool-seq data in eight separate years (Fig. 2). Frequencies at all shared and private SNPs were very highly correlated between replicates (R^2^ = 0.995 for years 1984, 1997, 2008), showing that the pool-seq data provided very precise information (Fig. S2). Frequency data from consecutive years also showed very consistent frequency estimates (Fig. 2). The cumulative frequency of the 6 mitotypes identified represented ~80% of the population. Other SNPs were identified in the dataset, but were not used as they could not be easily clustered or assigned to a mitotype due to the lack of important temporal frequency variation. Overall, the frequency pattern of the different mitotypes was remarkably stable, ruling out that the genetic composition of the mitochondrial population changed significantly over the study period. This therefore rules out that mitochondrial genetics explain the increased heat tolerance in the Vietnamese *Artemia* through time.

**Table S1.** Models used to generate effect sizes and variances for the meta-analysis of phenotypic adaptation (solid points, Fig 1A), which compared the overall temperature tolerance of VCH and SFB populations. Effect sizes are the log odds ratio of survival compared to SFB_84_. Where necessary, observation-level random effects were added to control for overdispersion. Notes: ^1^To estimate overall survival in this experiment, we fit models for survival in Phase 1 and Phase 2, then transformed, multiplied, and back-transformed the predicted survival rates. Standard errors were obtained by resampling. ^2^These models no longer have *Clutch* as a fixed term, so we control for possible variation in survival between the first and second clutches by including it here as a (non-dummy) random effect.

| Experiment | Fixed and random terms | $T_{SFB}$ | $T_{VCH}$ |
| --- | --- | --- | --- |
| | | Effect size ± SE | Effect size ± SE |
| Parental acclimation | $\sim VCH \text{ population} + (1\vert Family/Clutch^2)$ | VCH<sub>97</sub>: $-0.77 \pm 0.27$ | VCH<sub>97</sub>: $0.35 \pm 0.21$ |
| | | VCH<sub>08</sub>: $-0.30 \pm 0.27$ | VCH<sub>08</sub>: $1.18 \pm 0.19$ |
| Juvenile acclimation<sup>1</sup> | $\sim VCH \text{ population} + (1\vert Family/Observation)$ | VCH<sub>97</sub>: $0.38 \pm 0.68$ | VCH<sub>97</sub>: $0.19 \pm 0.46$ |
| | | VCH<sub>08</sub>: $-0.25 \pm 0.81$ | VCH<sub>08</sub>: $0.85 \pm 0.48$ |
| Microbiome | $\sim VCH \text{ population} + (1\vert Observation)$ | VCH<sub>97</sub>: $-0.18 \pm 0.28$ | VCH<sub>97</sub>: $0.52 \pm 0.23$ |
| | | VCH<sub>08</sub>: $-0.06 \pm 0.28$ | VCH<sub>08</sub>: $0.86 \pm 0.22$ |

**Table S2.** Significance of the tested effects for the individual experiments. Temp., temperature; treatm., treatment.

| Experiment | Fixed-effect term | T<sub>SFB</sub> |  | T<sub>VCH</sub> |  |
| --- | --- | --- | --- | --- | --- |
|  |  | Test statistic | P | Test statistic | P |
| **Additive genetic effects** | *Population* | $\chi^2_{(2)} = 1.7$ | 0.44 | $\chi^2_{(2)} = 3.6$ | 0.16 |
| **Juvenile acclimation** | *Population* | $\chi^2_{(2)} = 28.8$ | < 0.0001 | $\chi^2_{(2)} = 44.4$ | < 0.0001 |
| | *Temp. in Phase 1* | $\chi^2_{(1)} = 30.8$ | < 0.0001 | $\chi^2_{(1)} = 0.0$ | 0.91 |
| | *Population : Temp. in Phase 1* | $\chi^2_{(2)} = 2.3$ | 0.31 | $\chi^2_{(2)} = 1.0$ | 0.62 |
| **Parental acclimation** | *Clutch : Population* | $\chi^2_{(2)} = 2.1$ | 0.36 | $\chi^2_{(2)} = 1.8$ | 0.41 |
| | *Clutch : Parental treatm.* | $\chi^2_{(2)} = 3.3$ | 0.19 | $\chi^2_{(2)} = 3.8$ | 0.15 |
| | *Clutch : Population : Parental treatm.* | $\chi^2_{(4)} = 8.3$ | 0.08 | $\chi^2_{(4)} = 5.6$ | 0.23 |
| **Microbiome** | *Population* | - | - | - | - |
|  | *Microbiome* | - | - | - | - |
| | *Population : Microbiome* | $\chi^2_{(4)} = 15.7$ | 0.003 | $\chi^2_{(4)} = 18.4$ | 0.001 |

**Figure S1.**
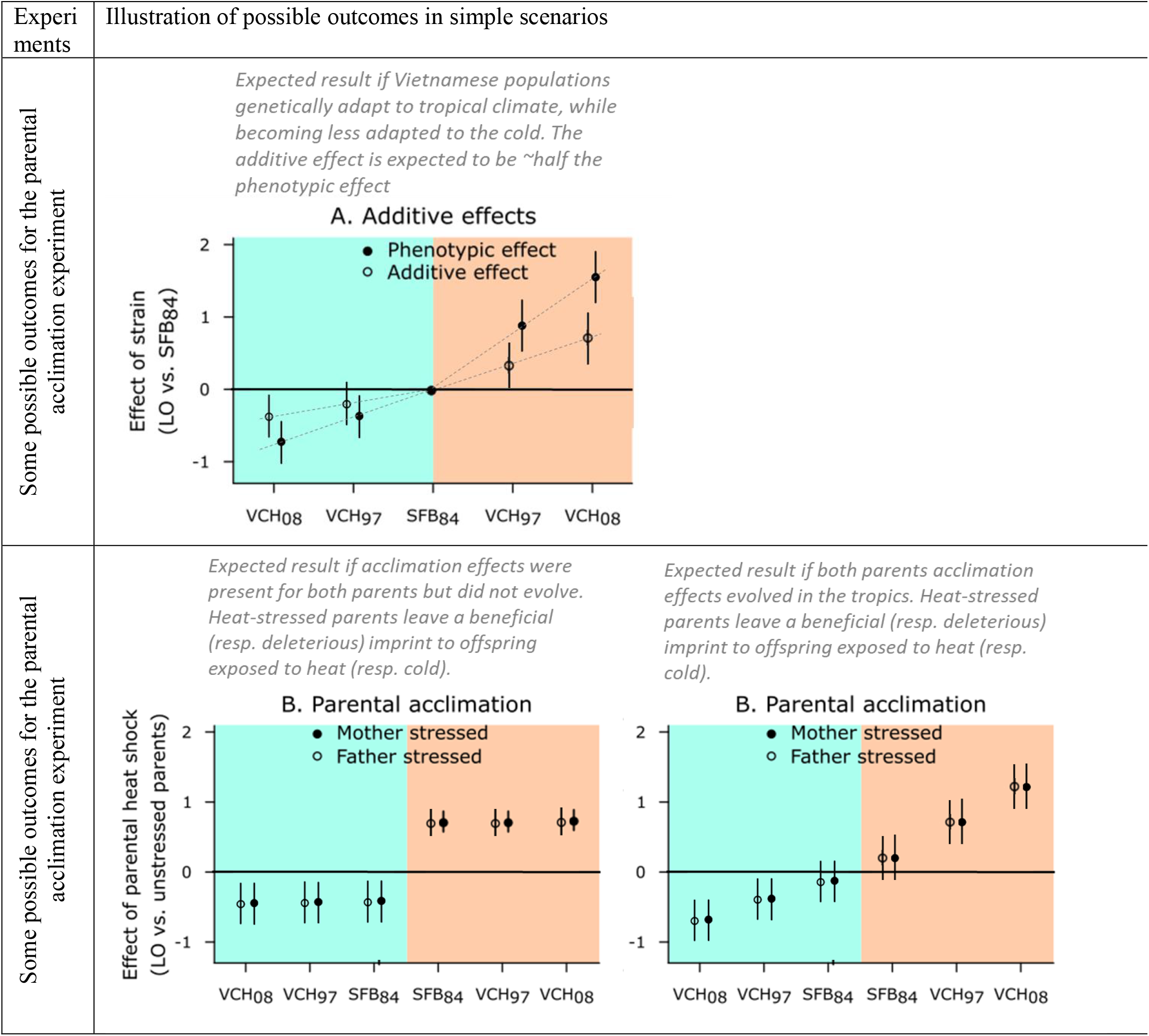
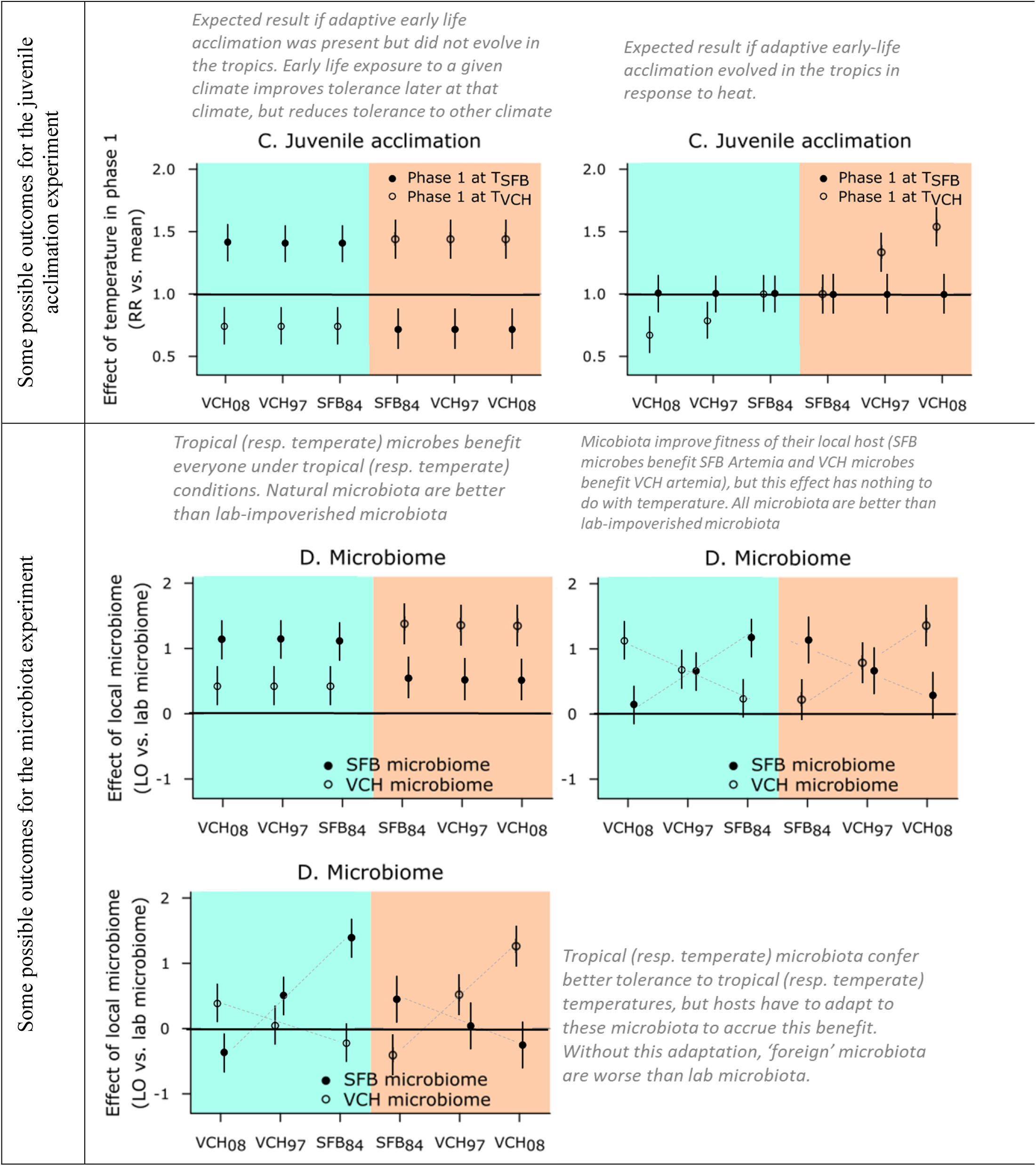
Illustration of possible outcomes for the different experiments, with simple scenarios described next to the figures.

**Figure S2.**
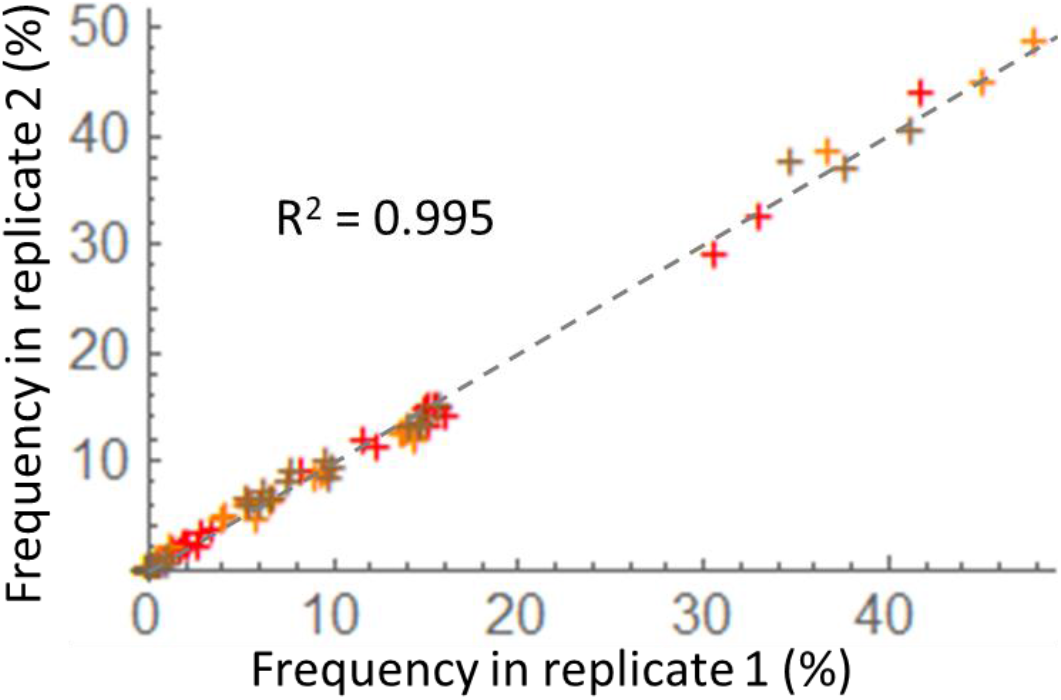
SNP frequency data quality. SNP frequency was independently estimated twice for years 1984 (red), 1997 (orange), and 2008 (brown). The figure reports the correlation between these replicated values for all SNPs used in Fig. 2 (all private and shared SNPs among sequenced individuals). R^2^ = 0.995 over all replicated measures. Dashed line is the 1:1 line.

**Figure S3.**
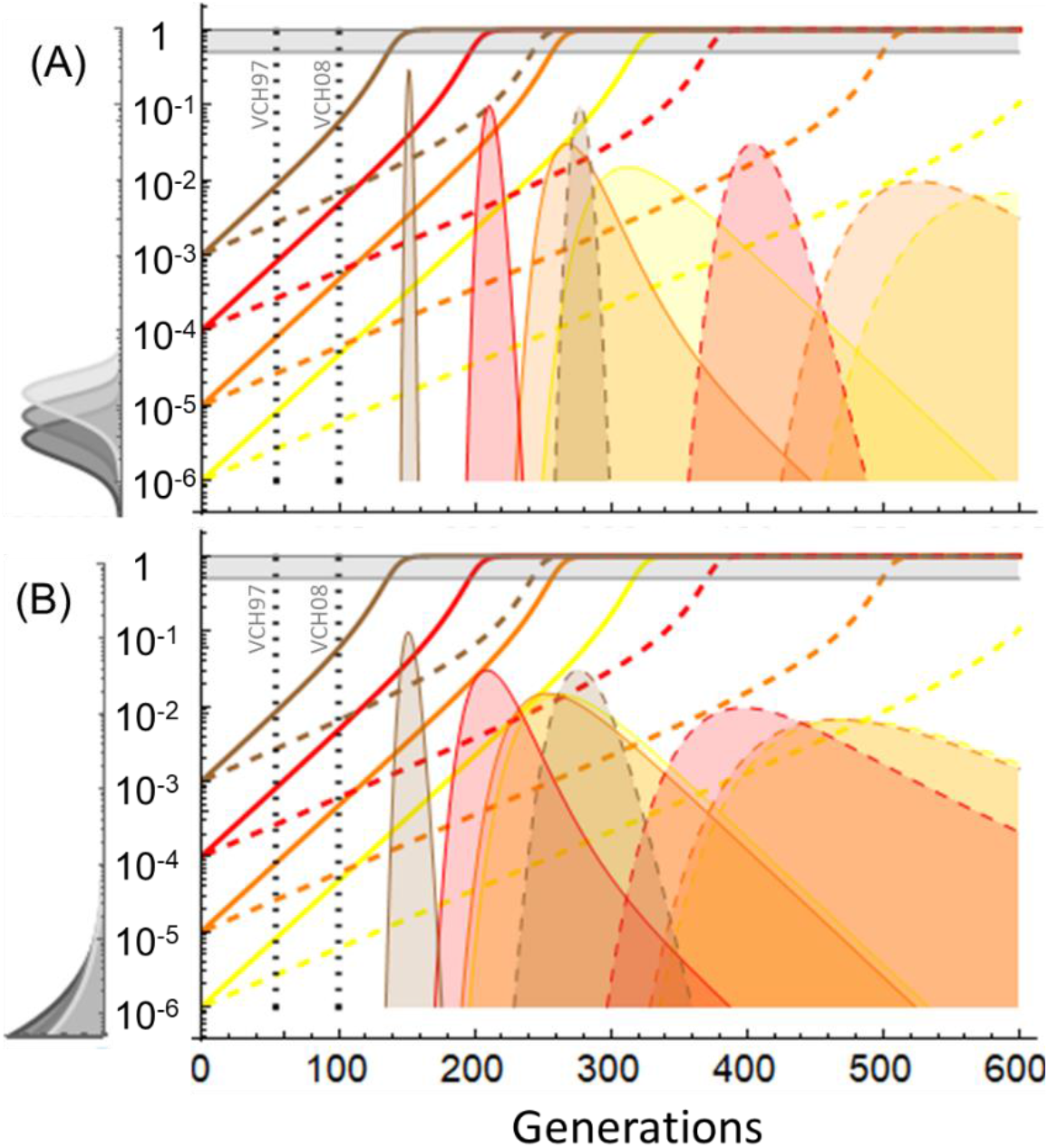
Frequency of a strongly beneficial recessive allele (*s* = 0.3) through time in a population of *N* = 10^7^ (panel A) or *N* = 10^6^ (panel B) individuals. Plain and dashed lines: exact deterministic frequency change for this beneficial allele given a dominance coefficient of *h* = 0.13 and *h* = 0.06, respectively. For these curves, the *x*-axis represents time in generations, and the *y*-axis is the frequency in log-scale. Different colors illustrate different initial frequencies, from 10^−6^ (yellow) to 10^−3^ (brown). For comparison, a newly arising mutant would start at 10^−7^ or 10^−6^ in panel A and B, respectively. The horizontal gray bar represents the [0.5,1] frequency range for the beneficial allele, within which it starts having a detectable effect on the mean fitness of the population (~10% increase in mean fitness). The vertical dotted lines correspond to our two dates of measurement (1997, at generation ~54, and 2008, at generation ~100). Colored surface areas show the corresponding probability density of half-sweep times (i.e. the number of generations to reach a frequency of ½) from a stochastic model with *N*_*e*_ = *N* (from Eq. 8 in (Martin and Lambert, 2015), same line and color code for dominance and initial frequency as for the deterministic curves). Here, the *x*-axis shows this half-sweep time and the *y*-axis the corresponding probability density. Distributions in grey to the left of the *y*-axis show the frequency distribution of the heat tolerance allele in its population of origin (SFB), at mutation (*u* = 10^−7^) selection balance, assuming that it reduces fitness at cold temperatures by a modest amount (*hs* = 0.02 dark grey, 0.01 grey or 0.005 light grey) compared to its strong advantage at high temperature. This Wright’s distribution is computed from Eq. 9.3.4 in (Crow and Kimura, 1970) with *N*_*e*_ = *N*. *y*-axis: frequency, *x*-axis: probability density (rescaled for readability on panel A by the inverse of the density at mean frequency). Overall, the figure shows that even in the most favorable conditions, adaptation caused by a recessive beneficial allele cannot be detected within the short 54 or 100 generations of our study. These are the most favorable conditions, as we consider (1) incompletely recessive beneficial alleles, with *h* = 0.13 corresponding to the point estimate based on observed survival in our experiment 1 for VCH_08_ which is certainly a maximum given that the point estimate in VCH_97_ would be *h* = 0 at most); (2) a very strong beneficial effect *s* = 0.3, comparable to the highest published field estimates of fitness effects in situations of intense selection pressures (e.g. insecticide resistance (Lenormand et al., 1999)); (3) plausible population sizes of *Artemia* populations, as census sizes in the field far exceed 10^6^ in Vinh Chau saltern; (4) a modest deleterious fitness effect of these beneficial alleles at cold temperature and standard mutation rate (10^−7^; a ten-fold larger mutation rate would not alter the conclusion, so that these mutations are unlikely to segregate at a higher frequency than 10^−3^ in the population of introduction). In addition, with such intense selection, beneficial alleles spread extremely fast when they reach a frequency above 0.2 (they then reach a frequency of 0.8 in ~20 generations), so that in this scenario, with a strongly beneficial allele we would be likely to observe a fitness change only between 1984 and 1997 or 1997 and 2008, but not both.

## Notes

### Competing Interest Statement

The authors have declared no competing interest.

